# EMMPRIN confers metabolic advantage for monocytes and macrophages to promote disease in a model of multiple sclerosis

**DOI:** 10.1101/2024.08.11.607460

**Authors:** DK Kaushik, A Das, C Silva, C D’Mello, LGN de Almeida, N Ghasemi, P Neri, A Dufour, NJ Bahlis, M Xue, VW Yong

**Affiliations:** Division of Biomedical Sciences, Faculty of Medicine, Memorial University of Newfoundland, St. John’s, Canada; Department of Clinical Neurosciences and Hotchkiss Brain Institute, Cumming School of Medicine, University of Calgary, Calgary, Canada; Departments of Biochemistry and Molecular Biology, and Physiology and Pharmacology, and McCaig Institute for Bone and Joint Health, Cumming School of Medicine, University of Calgary, Calgary, Canada; Department of Oncology and Arnie Charbonneau Cancer Institute, Cumming School of Medicine, University of Calgary, Calgary, Canada; Department of Cerebrovascular Diseases, The Second Affiliated Hospital of Zhengzhou University, Zhengzhou, China

## Abstract

Monocytes and monocyte-derived macrophages have important roles in the initiation and progression of multiple sclerosis (MS). These cells undergo metabolic reprogramming to generate immunophenotypes that promote leukocyte infiltration, axonal degeneration and demyelination, worsening MS pathology. The mechanisms that dictate metabolic programs in monocytes and macrophages in MS remain unclear. We previously reported that extracellular matrix metalloproteinase inducer (EMMPRIN, CD147), a glycoprotein that acts as a chaperone of monocarboxylate transporter 4 (MCT4), assisted with glycolysis-driven pro-inflammatory phenotype in macrophages in experimental autoimmune encephalomyelitis (EAE), an animal model of MS. Using newly-generated CCR2Cre^ERT2^:EMMPRIN^fl/fl^ (CCR2:EMMP) mice, we report that presymptomatic deletion of EMMPRIN in CCR2+ monocytes prevented or reduced clinical disability of EAE. This was correspondent with decreased infiltration of leukocytes into the CNS. Single cell RNA-seq of blood monocytes from EAE and proteomics analysis of macrophages from CCR2:EMMP^−/−^ mice revealed significant alterations in metabolic programs, particularly reduced glycolysis and elevated mitochondrial electron transport and fatty acid oxidation, which were linked to their reduced pro-inflammatory traits. Our findings implicate EMMPRIN as a key regulator of metabolic pathways that exacerbate pro-inflammatory functions of monocytes in MS.

## Introduction

Monocytes and monocyte-derived macrophages have critical roles in the initiation and progression of experimental autoimmune encephalomyelitis (EAE) and multiple sclerosis (MS) pathology^1–3^. Macrophages, during pro-inflammatory activation, undergo enhanced glycolysis^4^ which supports the production of reactive oxygen species (ROS) and inflammatory cytokines and chemokines^4–7^. Previously, we described that CNS-infiltrating macrophages in EAE and MS enhanced their expression of lactate dehydrogenase A (LDHA), a pyruvate-to-lactate converting enzyme, and hypoxia inducible factor-1α (HIF-1α), an upstream inducer of genes involved in glycolytic pathways^8, 9^. This led to the buildup of lactate within macrophages in the spinal cord of EAE mice^9^, export of which was facilitated by monocarboxylate transporter 4 (MCT4)^9^. We also found extracellular matrix metalloproteinase inducer (EMMPRIN, CD147), a transmembrane type-I glycoprotein^10, 11^, to be indispensable for chaperoning MCT4 to the plasma membrane of macrophages; preventing EMMPRIN-MCT4 interaction reduced macrophage transmigration *in vitro* and decreased the clinical severity of EAE mice^9^. Previous literature has shown that treatment with function-blocking antibody against EMMPRIN reduced the neuropathology of EAE^12^. Moreover, we observed elevated levels of soluble EMMPRIN in the cerebrospinal fluid (CSF) of individuals with secondary progressive multiple sclerosis (SPMS)^13^. These findings suggest a potential connection between EMMPRIN and the progression of MS, highlighting the clinical significance of targeting this glycoprotein.

Before they differentiate into CNS-infiltrated macrophages, monocytes must exit the bone marrow and migrate into secondary lymphoid tissues such as the spleen and lymph nodes. Monocytes that are Ly6C^hi^ CD11b^+^ CD45^+^ and Ly6G^−ve^ are pro-inflammatory and aggregate in large numbers within the perivascular spaces of cerebral post-capillary venules to form perivascular cuffs, as detected by CD45^+^or F4/80^+^ cells around laminin^+^ basement membranes^9, 14^, before entering the CNS parenchyma^1, 6, 9, 15^. CCR2 is the major chemokine receptor for CCL2 (also known as MCP-1) that homes monocytes to target tissues^6^. While EMMPRIN aids the infiltration of macrophages across the blood-brain barrier and into the CNS parenchyma^9^, its roles in dictating specific metabolic programs within CCR2^+^ monocytes during the early course of the disease is not known. Genetic constitutive knockout of EMMPRIN is lethal during embryogenesis in mice and is associated with developmental challenges in the few surviving offsprings^10, 16^. Therefore, to investigate the role of EMMPRIN specifically in monocytes and macrophages in adult-onset disease such as MS, we generated transgenic mice to inducibly and conditionally delete EMMPRIN from CCR2+ cells. With these mice, we now report the pivotal role of EMMPRIN in the maintenance of metabolic processes in pro-inflammatory monocytes and macrophages during EAE pathology. We show that EMMPRIN plays a crucial role in dictating metabolic pathways within CCR2+ monocytes and macrophages, aiding their proinflammatory functions and infiltration into the CNS to initiate EAE.

## RESULTS

### Manifestation of EAE is reduced in CCR2:EMMP^−/−^ mice

We generated EMMPRIN^fl/fl^ mice (Fig S1) and crossed them with CCR2Cre^ERT2^ line (gift from Dr. Becher^6^) to obtain a tamoxifen (tam)-inducible CCR2Cre^ERT2^:EMMPRIN^fl/fl^ line (CCR2:EMMP mice) (Fig 1A and Fig S1A-C). Confirmation of over 60% cre-recombinase activity was ascertained in CD11b+ Ly6C+ Ly6G-CD45+ monocytes by tdTomato expression in the red fluorescent channel when homozygous CCR2Cre^ERT2^ mice were crossed with homozygous Ai9 mice (Fig S2). Hematopoietic stem cells were then harvested from bone marrow of CCR2:EMMP mice and stimulated with 25ng/ml M-CSF (day 0, D0) followed by treatment with 5μM hydroxytamoxifen (4-OH) 24h later for the next 48h (day (D)1-D3); cells were harvested on D4 before they differentiated to macrophages. We confirmed the deletion of EMMPRIN in these monocytes using qRT-PCR (Fig 1B).

**Fig 1.**
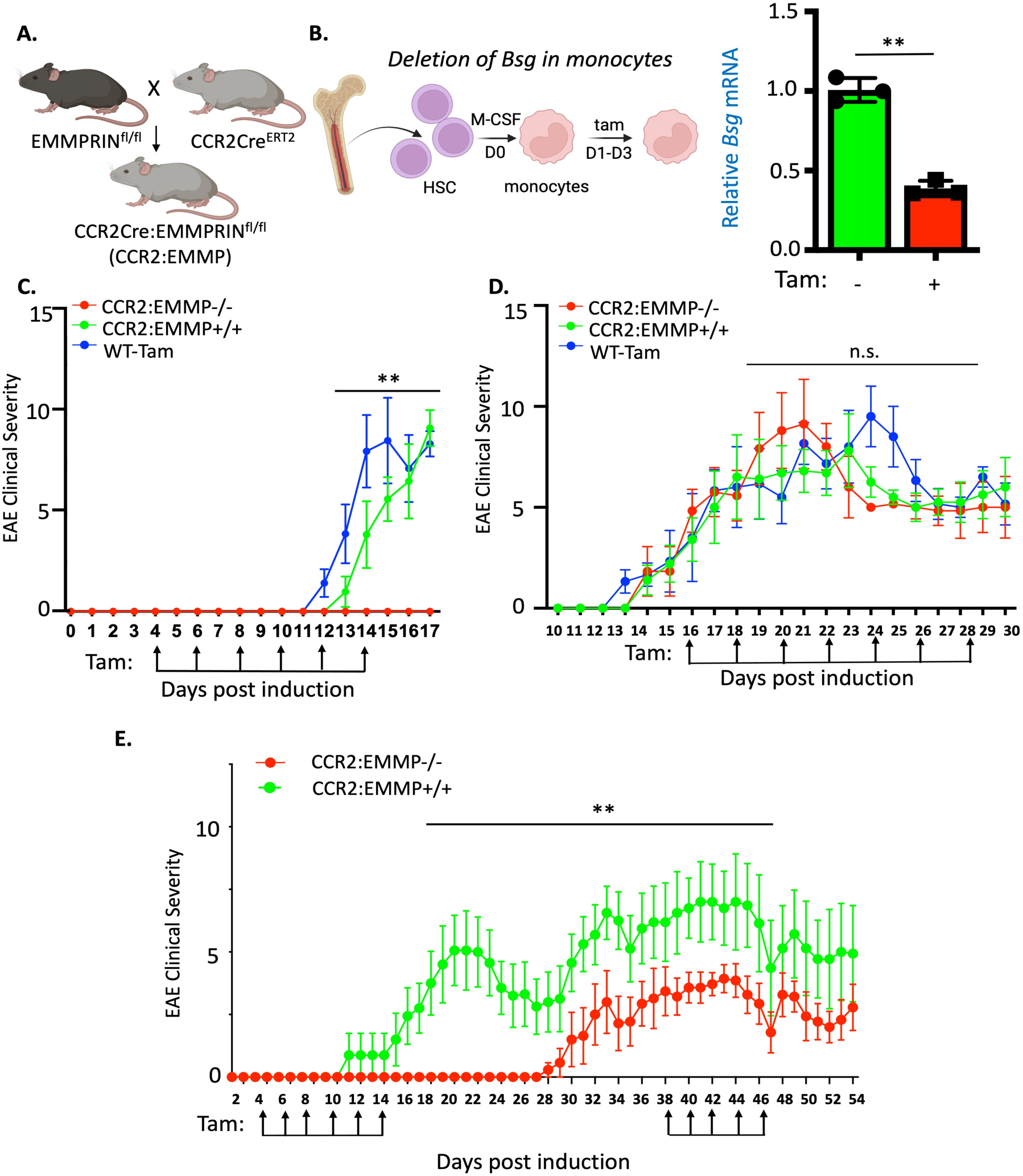
EMMPRIN deletion in monocytes in early EAE prevents or reduces clinical disability. **A)** Schematic showing the cross between EMMPRIN^fl/fl^ and CCR2Cre^ERT2^ lines to generate CCR2Cre:EMMPRIN^fl/fl^ mice (CCR2:EMMP). **B)** Monocytes harvested from the bone marrow of CCR2:EMMP mice were maintained in medium containing 25ng/ml M-CSF for a day before treating them with 5μM of 4-hydroxytamoxifen (tam) for the next two days. The cells were then harvested for RNA isolation and *Bsg* transcripts were assessed using qPCR. Student’s t-test, **p<0.01. **C)** EAE clinical severity on a 15-point scale showing no detectable disability in CCR2:EMMP mice that received 2mg/kg tamoxifen (CCR2:EMMP^−/−^) or those that received equal volume of corn oil (CCR2:EMMP^+/+^); tam injections started at D4 until D14 every two days. Wild-type (WT) mice injected with tam served as additional controls. Data were collected from at least 8 mice per experimental group. **D)** EAE disease course exhibiting severity in mice that received tam injections starting D16 until D28 and observed until D30. Data were collected from 6 mice per experimental group. **E)** Chronic EAE in CCR2:EMMP^+/+^ or CCR2:EMMP^−/−^ mice in a cohort similar to panel C but followed long-term until D54. Data were collected from 7 mice in the CCR2:EMMP^−/−^ and 8 mice in the CCR2:EMMP^+/+^ group. EAE data were analyzed using two-way ANOVA followed by Bonferroni post-hoc test. ** p<0.01. Graphs plotted as mean ± SD.

Circulating monocytes turnover about every 2 days. Therefore, to investigate the role of EMMPRIN in monocytes in EAE, we immunized (on day 0) CCR2:EMMP mice for the disease and injected tam every 2 days starting day D4 until D16 post-EAE (CCR2:EMMP^−/−^). When compared with wild-type (WT) mice receiving tam (WT-tam group) and littermate controls without tam (CCR2:EMMP^+/+^), where the EAE onset of clinical disability (impairment of tail functions) were on D11 and D12 respectively, there was no clinical manifestation of EAE in the CCR2:EMMP^−/−^ group even when the control mice were at peak disability (D17, paresis/paralysis of tail and all 4 limbs) (Fig 1C).

Next, we tested whether deletion of EMMPRIN post-onset of clinical signs, when monocytes have migrated into the CNS and become monocyte-derived macrophages^17^, would decrease the disability severity of EAE. We injected tam starting D16 post-EAE every two days until D28 (Fig 1D), but did not observe reduction of established disability of EAE in CCR2:EMMP^−/−^ group when compared with the controls (Fig 1D).

Another cohort of CCR2:EMMP mice that received early tam injections (D4-D14) was followed long-term and these did not exhibit clinical signs of EAE until D28 post-induction, a significant delay of approximately two weeks since the last tam injection at D14 (Fig 1E). Thereafter, clinical disease severity remained significantly milder throughout the observation period in these mice compared to the CCR2:EMMP^+/+^ cohort (Fig 1E). When we repeated tam injections starting D38 until D46 in CCR2:EMMP^−/−^ mice, there was no further change in the already milder disease course (Fig 1E).

Overall, these results suggest that EMMPRIN is important for early monocyte activity in EAE. Once disease is established with the many immune alterations that occur in EAE^18^, the role of EMMPRIN in monocytes and macrophages becomes less evident.

### EMMPRIN is required for monocyte trafficking into the CNS in EAE

We focused on the infiltration of immune cells into the CNS in EAE. CCR2:EMMP^−/−^ mice (from Fig 1C at tissue harvest on D18) had no obvious perivascular cuffs in the cerebellar white matter compared to readily observable cuffs in CCR2:EMMP^+/+^ mice (Fig 2A and 2B). We performed flow cytometry from spinal cords of mice with peak clinical disability (D18 in this cohort) and gated on CD45^hi^ Ly6G-CD11b+ population (Fig 2C). The results corroborate the lack of infiltration of EMMPRIN+ CCR2+ CD11b+ monocytes into the CNS in CCR2:EMMP^−/−^ mice, corresponding to the lack of clinical signs of EAE at this time period (Fig 1C). These effects were specific to CCR2:EMMP^−/−^ mice since the WT-tam group did not exhibit such differences in this myeloid population.

**Fig 2.**
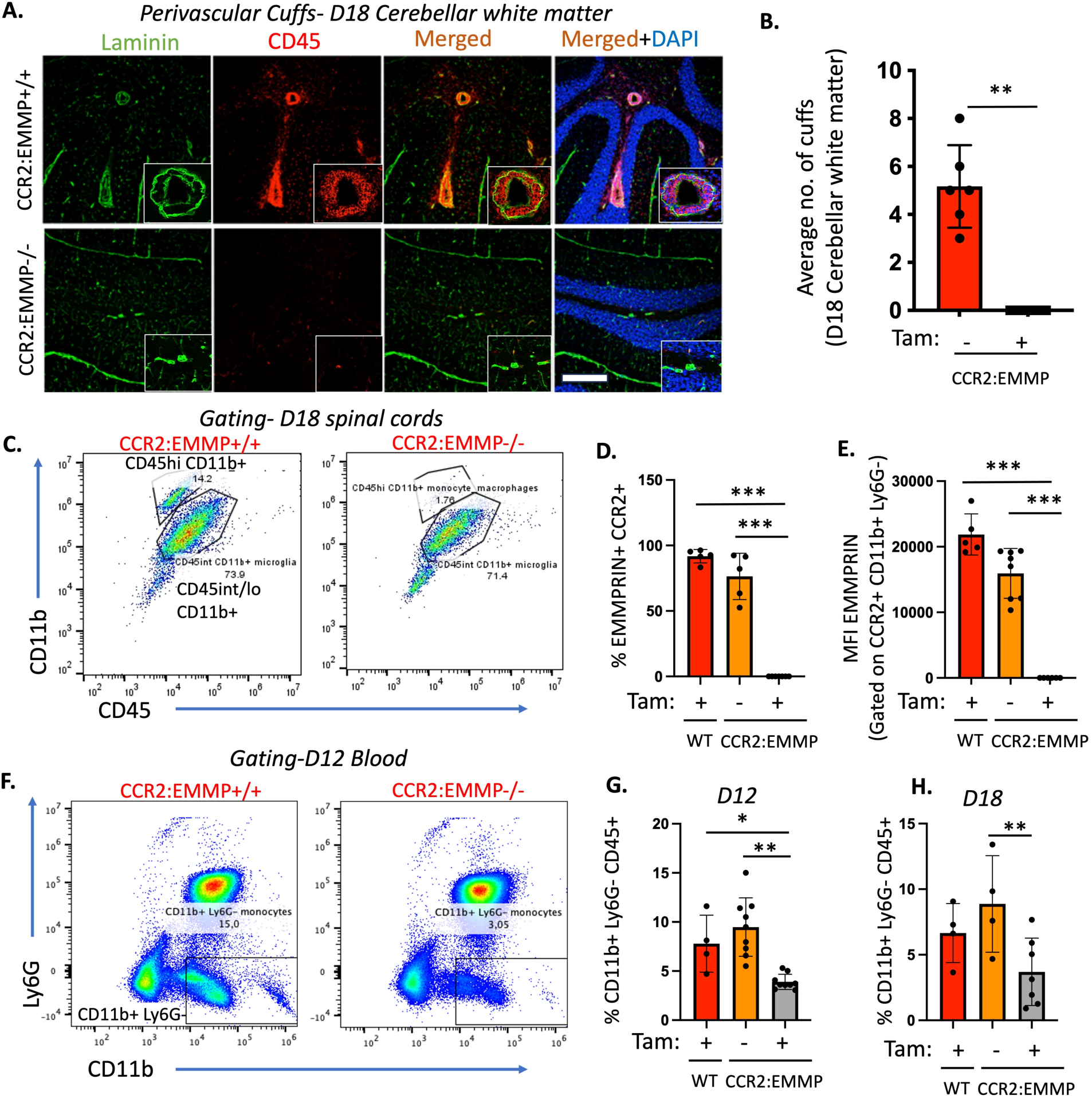
Analysis of CNS tissue for infiltration of monocyte-derived macrophages in CCR2:EMMP^−/−^ mice. **A)** Confocal images of perivascular cuffs in the cerebellar white matter of CCR2:EMMP^+/+^ and CCR2:EMMP^−/−^ D18 EAE showing pan-leukocytic marker, CD45 (red) encased in basement membrane delineated by laminin staining (green). Insets showing enlarged cuffs with CD45+ cells within. Scale-50μm. **B)** Histograms denoting average number of cuffs observed in the two groups. Data analyzed by student’s T-test. **p<0.05, N of 6 per group. **C)** Spinal cords were harvested from CCR2:EMMP^+/+^ and CCR2:EMMP^−/−^ mice at D18 after immunization and subjected to flow cytometry: Singlet viable cells were gated on CD11b and CD45 for CD45hi CD11b+ monocyte/macrophages as shown in dot-plot for CCR2:EMMP^−/−^ and CCR2:EMMP^+/+^ examples; there were few monocyte/macrophages in the spinal cord of CCR2:EMMP^−/−^ mice. **D)** Analysis of % EMMPRIN+ CCR2+ infiltrated macrophages as well as **E)** expression of EMMPRIN levels (MFI = mean fluorescence intensity) in CCR2+ CD11b+LY6G-cells. **F)** Dot plot exhibiting Ly6G and CD11b staining from blood of CCR2:EMMP^+/+^ and CCR2:EMMP^−/−^ D12 EAE mice which is quantified as %CD11b+ Ly6G-CD45+ cells in **(G)**. Histograms showing %CD11b+ Ly6G-CD45+ cells from D18 EAE in WT, CCR2:EMMP^+/+^ and CCR2:EMMP^−/−^ mice. Flow plots shown in D, E, G and H were compared using one-way ANOVA with Bonferroni post-hoc test. *p<0.05, **p<0.01 and ***p<0.001. Data represented as mean ± SD.

We investigated whether the lack of CNS infiltration in CCR2:EMMP^−/−^ mice was due to reduction in the number of circulating monocytes during onset and peak EAE. We enumerated proinflammatory monocytes in blood and while there were no significant differences in CD11b+ Ly6G-CD45+ cells in WT-tam and CCR2:EMMP^+/+^ mice, their numbers significantly decreased in D12 and D18 post-immunization in CCR2:EMMP^−/−^ blood (Fig 2F-H).

These observations provide evidence that EMMPRIN knockout from monocytes renders these cells with reduced CNS-infiltration capabilities. It is important to note that migration of Ly6C+ CD11b+ Ly6G-cells from bone marrow to blood was not affected in the non-EAE healthy CCR2:EMMP^−/−^ mice (Supplemental Fig 3B). This suggests that knocking down EMMPRIN from monocytes in healthy mice does not impact their bone marrow egress and likely will not impact the homeostatic functions of monocytes.

### Defining the distinct monocyte clusters and subclusters during pre-onset EAE in CCR2:EMMP^+/+^ and CCR2:EMMP^−/−^ mice

Given that EMMPRIN knockout from monocytes renders these cells with reduced CNS-infiltration capacity, we sought to understand the mechanisms behind EMMPRIN’s role in monocytes during early stages of EAE. We sorted CD11b^+^ immune cells from three D7 EAE CCR2:EMMP^−/−^ mice and three D7 CCR2:EMMP^+/+^ mice using the EasySep CD11b^+^ magnetic sorting kit (StemCell Technologies) and performed single cell RNA-sequencing (sc-RNA seq) using 10X Genomics Chromium platform followed by NovaSeq Illumina sequencing. The R package Seurat was used to perform quality control, principal component analysis and clustering^19^. Post quality control, the expression matrix contained a total of 20,306 cells (n= 9,195 cells from CCR2:EMMP^+/+^ and n=11,111 cells from CCR2:EMMP^−/−^ groups) with 31,527 features/genes. Unsupervised clustering of total cells from CCR2:EMMP^−/−^ and CCR2:EMMP^+/+^ mice with EAE, based on 2000 variable genes and 30 significant principal components, delineated 14 clusters. Clusters were annotated using canonical markers which defined 12 neutrophil clusters (Clusters 0-7, 9 and 11), a B-cell cluster (Cluster 10), an NK-cell cluster (Cluster 12), a cluster of cells with unknown phenotype (Cluster 13), and a pro-inflammatory monocyte cluster (Cluster 8; marked as CD45^+^ CD11b^+^ Ly6C^hi^ Ly6G^−ve^) (Fig 3A). Cluster 8 was defined by high levels of CCR2 and Ly6C2 expression (Fig 3B and 3C).

**Fig 3.**
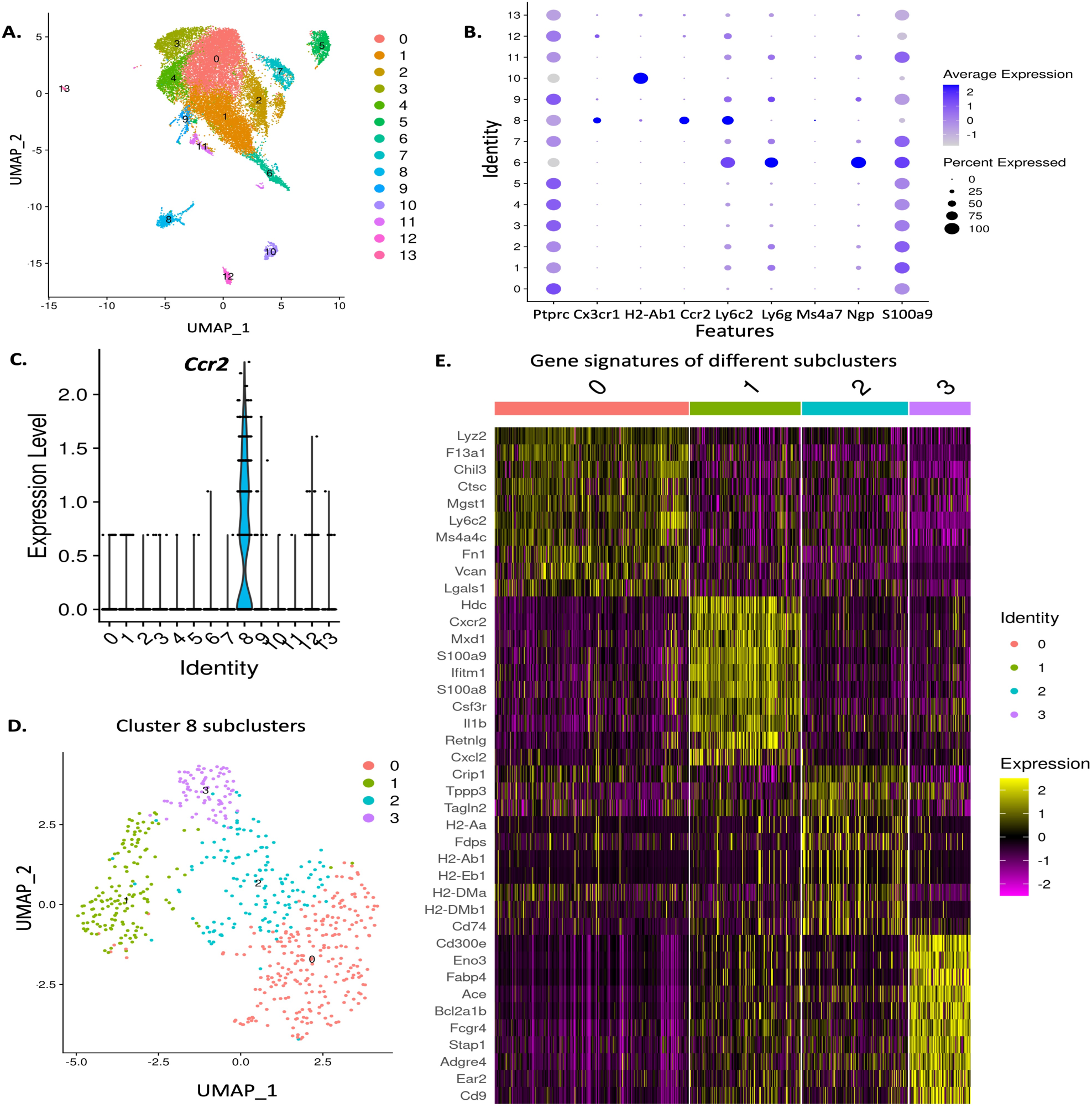
scRNAseq using 10X genomics on CD11b+ cells from D7 EAE blood in CCR2:EMMP^+/+^ and CCR2:EMMP^−/−^ mice. **A)** UMAP Projection of CD11b+ cells isolated using CD11b+ positive selection magnetic sorting from blood of 3 CCR2:EMMP^+/+^ and 3 CCR2:EMMP^−/−^ mice at D7 post-immunization. Clusters 0-7, 9 and 11 represent neutrophils, cluster 10 represents B cells, cluster 12 denotes NK-cells and cluster 13 for unknown cells. Cluster 8 represents the pro-inflammatory monocyte cluster as corroborated by high levels of CCR2 and Ly6C2 expression **(B-C)**. **D)** UMAP projection of Cluster 8 within the blood of CCR2:EMMP^−/−^ and CCR2:EMMP^+/+^ mice revealed four distinct subsets of monocytes that were identified as- 0, 1, 2 and 3. **E)** Heat map analysis of the top 10 DEGs among the different subclusters coloured for their expression levels.

To further assess subsets within the Cluster 8 population, this cluster was separated from the other cells in the dataset and re-clustered. Clustering analysis using 20 principal components and 0.5 resolution revealed four distinct subsets of monocytes that were identified as subclusters 0, 1, 2 and 3 (Fig 3D). A heat map of the top 10 differentially expressed genes (DEGs) among the different clusters revealed subcluster 0 to be CCR2^high^ Ly6C^high^ Cx3CR1^low^ which is enriched in glutathione pathway as it had high levels of *Mgst1* transcripts^20^ (Fig 3E). This subcluster also exhibited enrichment in genes encoding extracellular matrix proteins, versican (*Vcan*) and fibronectin (*Fn1*). Subcluster 1 expressed high levels of *Il-1β*, chemokine *Cxcl2,* and its receptor, *Cxcr2* suggesting this subcluster to be associated with EAE pathology and inflammation in general^21^. Subcluster 2 had the highest expression of HLA class II genes including *H2-Aa*, *H2-Ab1*, *H2-Eb1*, *H2-DMa* and *H2-DMb1* suggesting their role in antigen presentation and involvement in initiating EAE pathology^22^ (Fig 3E). Characterization of the DEGs defining subcluster 2 using gene set enrichment analysis tools such as EnrichR (https://www.maayanlab.cloud/Enrichr/)^23^ and Ingenuity Pathway Analysis (IPA, Qiagen) revealed this subcluster to engage in cholesterol and fatty acid metabolism and mTORC1 signaling (Supplemental Fig 4B and 4C). Subcluster 3 expressed high levels of *Fcgr4,* a low affinity immunoglobulin gamma Fc-receptor gene, as well as *Fabp4*, an intracellular lipid chaperone (Fig 3E). Overall, we identified key characteristics of distinct monocyte subpopulations during the pre-onset EAE among the CCR2:EMMP^−/−^ and CCR2:EMMP^+/+^ groups.

### EMMPRIN deletion reduces monocytes in EAE blood while altering their metabolism and pro-inflammatory genes

UMAP clustering showing a distribution of CD11b+ cells among the three CCR2:EMMP^−/−^ and three CCR2:EMMP^+/+^ mice at D7 post-immunization revealed no major differences in cluster 0-7, 9, 10, 11, 12 and 13 of CCR2:EMMP^−/−^ mice (Fig 4A). However, CCR2:EMMP^−/−^ mice had a significant reduction in the percentage of monocytes in Cluster 8 when compared to the control group (Fig 4A and Fig 4B). Proportional analysis (Fig 4C) and tSNE clustering (Fig 4D) confirmed decrease in Cluster 8 monocytes compared to other CD11b+ populations in CCR2:EMMP^−/−^ mice. Importantly, a feature plot showing expression of the *Bsg* gene (which encodes EMMPRIN) across all clusters confirmed reduction in *Bsg* expression in the remaining cells within Cluster 8 in CCR2:EMMP^−/−^ mice (Fig 4E). Comparison of top 20 DEGs within Cluster 8 revealed notable reduction in genes related to mitochondrial respiration (*mt-Cytb*, *Ndufa3*, *Mt-Nd3)* and inflammatory pathways (*Mt1*, *ccl9*, *Ifi30*) in CCR2:EMMP^−/−^ group. On the other hand, *Slfn4*, a gene which is associated with myelopoiesis^24^ and involved in inducing myeloid derived suppressor cells^25^, was elevated in monocytes from CCR2:EMMP^−/−^ mice suggesting the possibility of monocytes failing to differentiate to macrophages (Fig 4F). This was further supported by an elevation in *Csf3r*, a granulocyte colony stimulating factor receptor which is known to be enriched in the precursor cells in the bone marrow^26^. Moreover, genes related to the interferon pathway with known anti-inflammatory functions (*Oasl2, Ifit3, and Ifit1*) were found to be enriched in CCR2:EMMP^−/−^ monocytes (Fig 4F).

**Fig 4.**
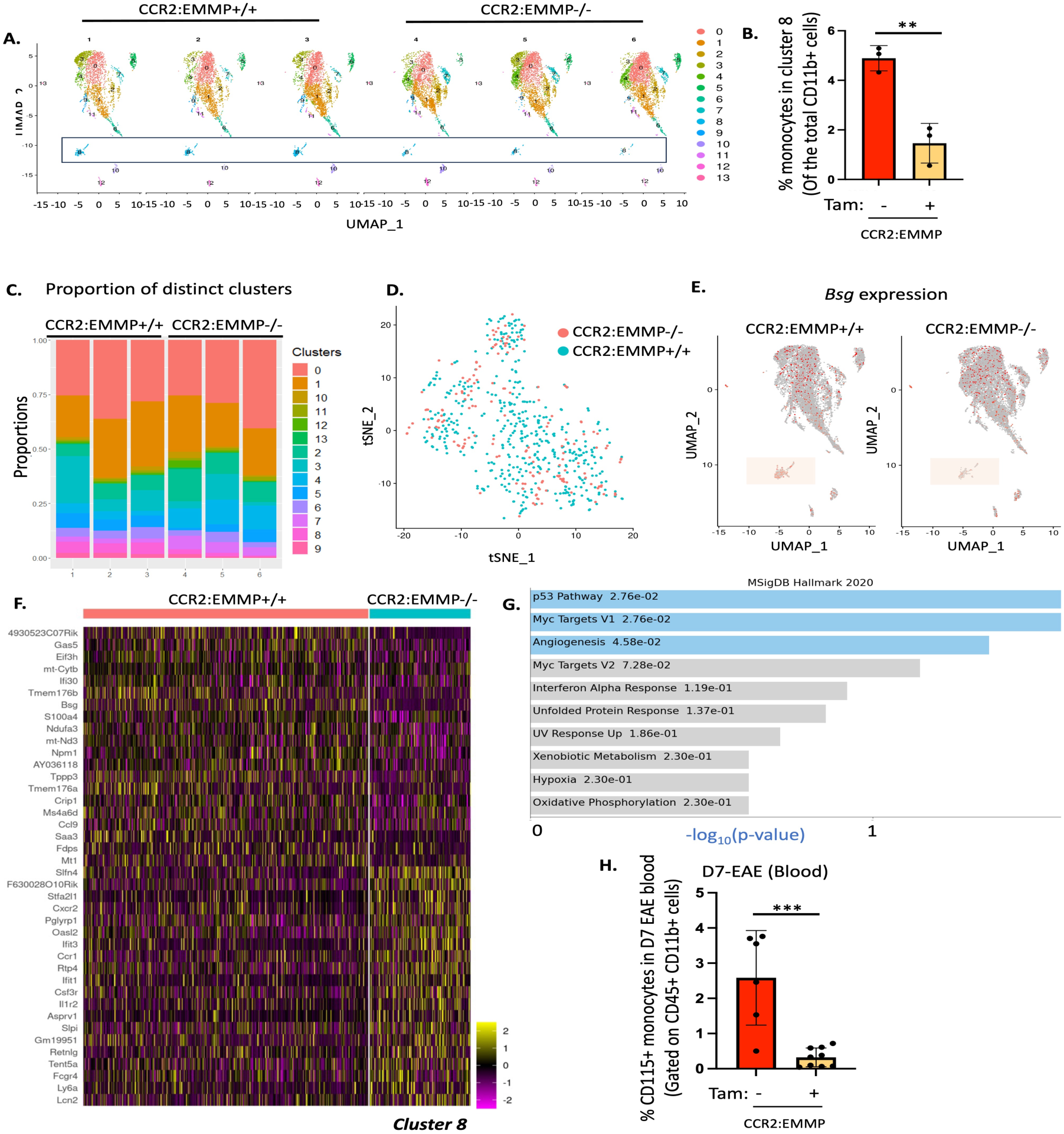
Effect of EMMPRIN knockdown in CCR2+ monocytes. **A)** UMAP projections from 3 CCR2:EMMP^+/+^ and 3 CCR2:EMMP^−/−^ mice post-immunization (D7) for EAE shown separately for effects on Cluster 8 and quantified as % monocytes of the total CD11b+ cells in **(B). C)** Graph showing proportion of different clusters among the CCR2:EMMP^+/+^ and CCR2:EMMP^−/−^ groups. **D)** tSNE representation showing superimposed Cluster 8 to denote spatial representation of this cluster in the two groups. **E)** UMAP projection highlighting cells for *Bsg* transcriptomic expression in all the clusters with emphasis on Cluster 8. **F)** Heat map showing top 20 DEGs downregulated (top) or 20 DEGs upregulated (bottom) within Cluster 8 of 3 CCR2:EMMP^+/+^ and 3 CCR2:EMMP^−/−^ mice. Expression levels of genes are colour-coded for each cell. **G)** MsiDB Hallmark 2020 database analysis of the major pathways that change in Cluster 8. **H)** Flow cytometry analysis of %CD115+ monocytes in D7 EAE blood gated on CD45+ CD11b+ Ly6G-viable cells. Datasets compared using student’s t-test; ***p<0.001. Data represented as mean± SD.

CCR2:EMMP^−/−^ monocytes also increased their expression of *Il-1r2*, a non-signaling receptor for IL-1β which prevents the activity of IL-1β by serving as a decoy receptor to IL-1R^27^. We also observed an increase in *Fcgr4* (Fc-receptor gamma gene) in CCR2:EMMP^−/−^ monocytes, suggesting the possibility of higher opsonin-dependent phagocytosis^28^ by these monocytes. Since Fc-receptor internalization is dictated by phagocytic process, which in-turn is driven by OXPHOS, increase in Fc-receptor transcripts signifies that CCR2:EMMP^−/−^ monocytes bear homeostatic traits while maintaining intact immune-competency. Further, CCR2:EMMP^−/−^ monocytes also increased their expression of CCR1, a chemokine receptor to chemokines including CCL1, which is known to induce a Th2 response ^29^. Using the MSigDB Hallmark 2020 database^30^, we identified multiple pathways belonging to ‘angiogenesis’, ‘hypoxia’ and ‘oxidative phosphorylation’ enriched in Cluster 8 in CCR2:EMMP^−/−^ mice (Fig 4G). Considering EMMPRIN’s established role in inducing matrix metalloproteinases (MMPs)^31, 32^, it was surprising to note that transcripts of genes pertaining to MMP pathway did not feature within the top 20 DEGs.

Heat map showed significant increase in *Slfn4* transcripts in Cluster 8 of CCR2:EMMP^−/−^ mice (Fig 4F). *Slfn4* is a negative regulator of differentiation of monocytes to macrophages *in vitro*^24^, confirming the reduction in differentiation efficiency in CCR2:EMMP^−/−^ monocytes. To test this possibility, we performed flow-cytometry of CD115+ (CSF-1R+) monocytes and found them to be significantly reduced in the CCR2:EMMP^−/−^ group (Fig 4H).

The monocyte subclusters, 0, 1, 2 and 3 were all decreased in the CCR2:EMMP^−/−^ group with the strongest effect seen for subcluster 2 (Fig 5 and supplemental Fig 4A). Further, loss of EMMPRIN from the subclusters of Cluster 8 in CCR2:EMMP^−/−^ mice validated the knockdown of this glycoprotein in different monocyte sub-populations (Fig 5B). All these subclusters expressed CD68 (endo-lysosome/phagocytic marker) and CSF1R (CD115), a receptor for CSF-1 (M-CSF), and a signaling pathway important for macrophage differentiation (Fig 5C and 5D). However, only three of the four subclusters, i.e. 0, 1 and 2, expressed high levels of CCR2 and Ly6C2 (Fig 5E and 5F).

**Fig 5.**
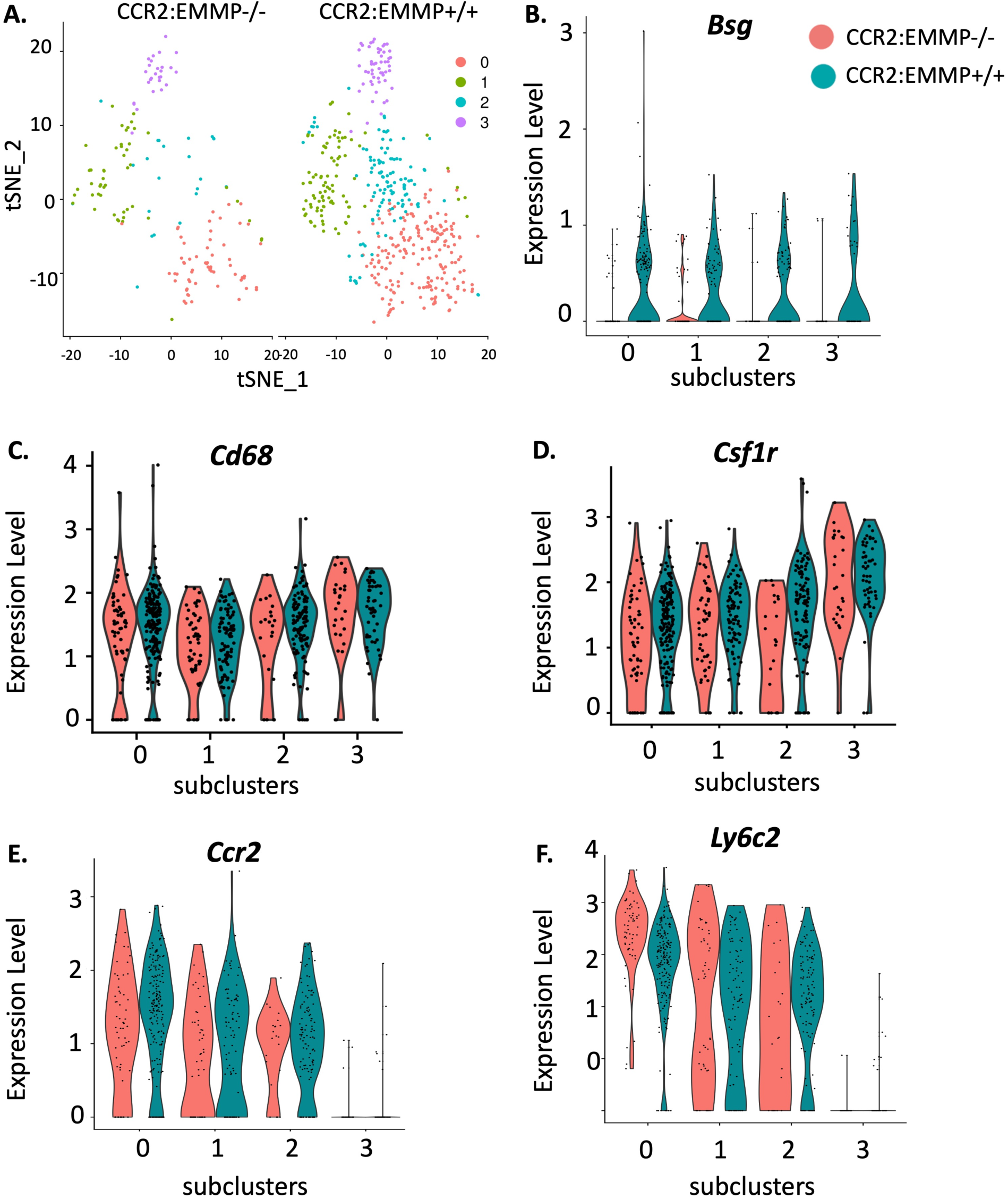
Effects of different canonical markers of individual subclusters in response to EMMPRIN knockdown. **A)** tSNE representation of subclusters 0, 1, 2 and 3 comparing between the CCR2:EMMP^+/+^ and CCR2:EMMP^−/−^ groups. **B)** Violin plot exhibiting transcriptomic levels of *Bsg* in individual subclusters confirming deletion of this gene in the subclusters. **C-D)** Expression levels of *Cd68* and *Csf1r* transcripts in CCR2:EMMP^+/+^ and CCR2:EMMP^−/−^ groups. **D-E)** Expression levels of *Ccr2* and *Ly6c2* transcripts in all the subclusters in CCR2:EMMP^+/+^ and CCR2:EMMP^−/−^ groups.

These findings strongly support EMMPRIN’s role in sustaining essential metabolic needs in pro-inflammatory monocytes and contributing to their pro-inflammatory phenotype. These findings also inform on the complexity of metabolic pathways that range from oxidative phosphorylation to glucose metabolism that enable monocyte activation during the early course of EAE pathology.

### EMMPRIN supports pro-inflammatory functions of macrophages

To identify how EMMPRIN impacts differentiated macrophages, we harvested bone marrow monocytes from CCR2:EMMP mice and differentiated them for 7 days to generate bone marrow-derived macrophages (BMDMs). We induced EMMPRIN deletion with 24h treatment of 5µM 4-OH tam before stimulating with 100ng/ml LPS for an additional 24h (Fig 6A). Tam-alone group served as control for tam-induced changes. Live/dead assay revealed no effects of tam on either cell proliferation or cell death in BMDMs (Supplemental Fig 5A). While LPS stimulation enhanced EMMPRIN expression in BMDMs, 4-OH tam treatment efficiently reduced the levels of EMMPRIN as confirmed by Western blotting (Fig 6A). Further, this led to reduction in the transcripts of hexokinase 1 (*hk1*), a rate-limiting glycolytic enzyme (Fig 6B), and reduction of *slc16a3* transcripts, the MCT-4 gene (Fig 6C). These findings confirm a reduction in the glycolytic processes in CCR2:EMMP^−/−^ BMDMs. In support, significant reduction in lactate levels was observed within the supernatants of CCR2:EMMP^−/−^ BMDMs (LPS+ 4-OH group), while treatment with tam in WT BMDMs did not affect lactate secretion in response to LPS stimulation (Fig 6D).

**Fig 6.**
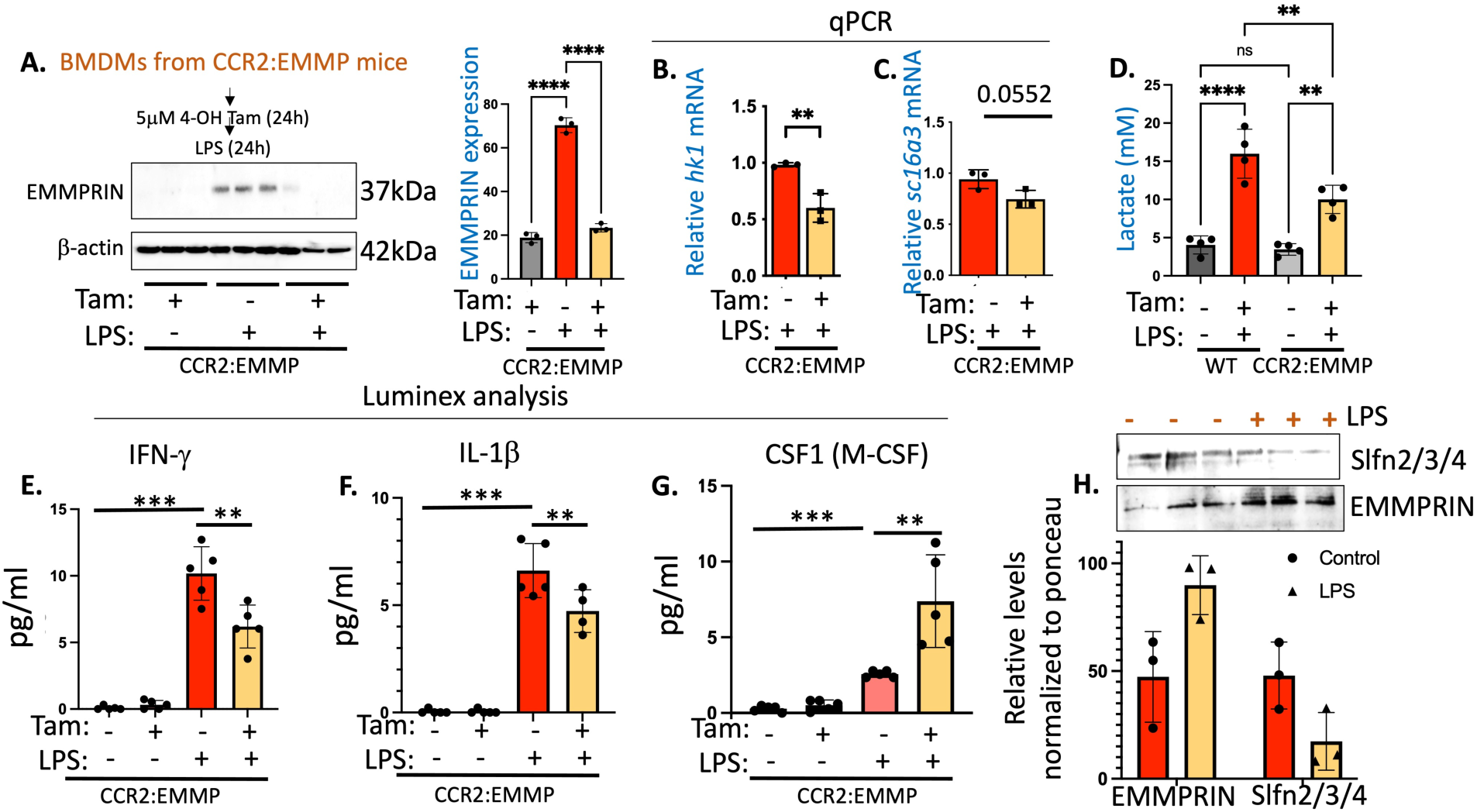
EMMPRIN deletion in differentiated macrophages. **A**) Bone marrow derived macrophages (BMDMs) were harvested from CCR2:EMMP mice and were differentiated for 7 days in M-CSF media before treating with 5μM 4-OH tamoxifen (Tam) for 24h prior to 24h LPS treatment. Western blot and its quantification (normalized to β-actin) show the magnitude of EMMPRIN deletion. Groups compared using one-way ANOVA. **B)** qPCR analysis of hexokinase (*hk1*) transcripts in LPS+tam condition when compared with LPS-alone treatment condition. **C)** qPCR analysis of MCT-4 (*Slc16a3*) transcript levels. Groups compared using student’s t-test. **D)** Lactate assay of the supernatants of BMDMs from WT and CCR2:EMMP mice treated with or without 5μM tam. **E-F)** Luminex analysis from the supernatants of BMDMs in different treatment conditions assessed for IFN-γ (E), IL-1β (F) and CSF-1 (M-CSF). **G**) Groups compared using one-way ANOVA. **H)** Western blot assessment for Slfn2/3/4 and EMMPRIN expression (normalized to ponceau) in response to LPS treatment. *p<0.05, **p<0.01, ***p<0.001. Data represented as ± SD.

Enhanced glycolysis is associated with the production of pro-inflammatory cyto-chemokines^8, 9^, therefore we performed cytokine analysis from the supernatants of CCR2:EMMP BMDMs. We found CCR2:EMMP^−/−^ BMDMs to exhibit significant reduction in LPS-induced interferon gamma (IFNγ) and IL-1β levels over LPS-alone group (Fig 6E and 6F). To corroborate reduction in CD115+ CD11b+ monocytes from CCR2:EMMP^−/−^ D7 EAE in CCR2:EMMP^−/−^ blood (Fig 4J), we measured the levels of CSF-1 (M-CSF, ligand for CSF-1R or CD115) to address whether EMMPRIN deletion affected the differentiation program of CCR2:EMMP^−/−^ BMDMs. Intriguingly, CSF1 levels were significantly elevated in LPS+tam group over LPS-alone group (Fig 6G), which is potentially a result of compensation to overcome the reduced differentiation of CCR2:EMMP^−/−^ BMDMs. To confirm if Slfn4, an ‘anti-differentiation’ marker observed to be high in CCR2:EMMP^−/−^ monocyte Cluster 8, decreases with inflammation, we performed Western blotting for Slfn2/3/4 in response to LPS in WT macrophages and found it to be decreased in LPS-treated group (Fig 6H); we also observed an inverse correlation between Slfn4 and EMMPRIN expression. Thus, it is likely that EMMPRIN deletion impairs differentiation of monocytes to macrophages, likely through metabolic perturbations during early course of EAE, a hypothesis which remains to be established.

### EMMPRIN dictates global metabolic changes in differentiated macrophages

To study changes in protein network upon EMMPRIN deletion, we performed unbiased analysis using quantitative proteomics analysis on the protein lysates from LPS-treated and LPS+ 4-OH-tam treated BMDMs (Fig 7A). 4-OH-tam treatment alone did not affect the proteome of these macrophages i.e. WT+tam proteome was not significantly different from that of WT-no tam proteome (Supplemental Fig 5B). EMMPRIN deletion led to significant changes in BMDM proteome predominantly affecting metabolism-related proteins (Fig 7A) which is confirmed by network analysis using STRING-DB (Fig 7B and 7C). In CCR2:EMMP^−/−^ BMDMs, we found significant reduction in Ndufs1, which codes for complex I subunit in the electron transport chain and potentially associated with pro-inflammatory functions including the generation of reactive oxygen species (ROS) as shown for Ndufs4 in myeloid cells^33^ (Fig 7A). We also observed decrease in expression levels of NLRP3, an inflammasome complex which drives IL-1β processing in these cells (Fig 7B), thus providing the basis of reduced IL-1β generation in CCR2:EMMP^−/−^ BMDMs. The metabolism-related proteins that were upregulated in response to EMMPRIN knockdown included acyl coenzyme A (acyl-CoA) thioesterase 7 (Acot7), an enzyme known to antagonize fatty acid utilization and breakdown^34,35^, mitochondrially encoded cytochrome c oxidase I (mtco1), a component of cytochrome c oxidase and negatively correlated with inflammation^36^, and Acaa2, an enzyme involved in the last step of fatty acid oxidation, and responsible for the generation of acetyl-CoA^37^. As noted in Fig 7B, pathways pertaining to aerobic glycolysis (BSG, phospofructokinase 1 (PFK1), HK1, HK2, PFKFB3, and SLC16a3) were among those that decreased significantly in response to EMMPRIN knockdown (Fig 7B). Gene pathway analysis using Metascape identified proteins pertaining to glycolysis were significantly downregulated in response to EMMPRIN knockdown (Fig 7B and 7C), while proteins belonging to mitochondrial electron transport chain along with fatty acid oxidation were enriched. Overall, these results highlight the prominent role of EMMPRIN in assisting distinct metabolic pathways during inflammatory processes within monocytes and macrophages.

**Fig 7.**
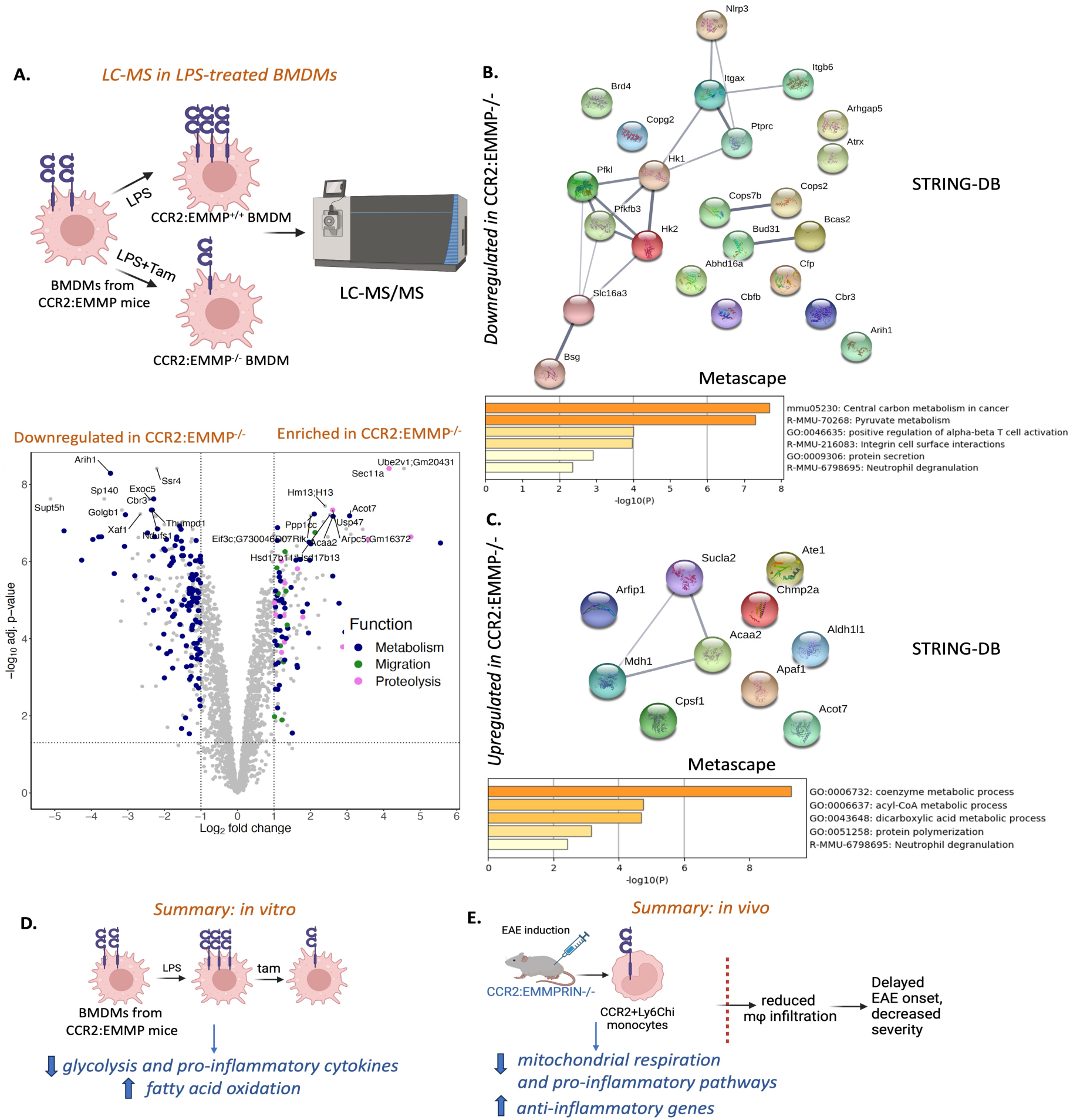
Liquid chromatography-mass spectrometry (LC-MS/MS) proteomics in LPS-treated BMDMs. **A)** Schematic showing treatment of BMDMs with LPS without tam or LPS treatment in conjunction with 5μM 4-OH tam prior to LC-MS/MS (upper panel). Volcano plot analysis of proteins enriched or downregulated in CCR2:EMMP−/− BMDMs (lower panel). Data points plotted as Log2 fold change vs –log10adj. p-value. Proteins pertaining to metabolism, migration or proteolysis that exhibit significant changes are highlighted. Most changes are observed in pathways pertaining to metabolism. **B-C)** Network analysis using STRING-DB and pathway analysis by Metascape depicting downregulated **(B)** and upregulated **(C)** protein networks. **D-E)** Schematic summary of the findings on the relevance of EMMPRIN in monocytes and macrophages from in-vitro **(D)** and in-vivo **(E)** experiments from this study.

## DISCUSSION

Circulating monocytes exhibit considerable differences in their metabolic program during relapse of a disease when compared with remission, as observed in the case of gout^38^. Some of the pathways that support inflammatory programs in classical monocytes (CD14+CD16-) in gout patients include arachidonic acid metabolism while non-classical homeostatic monocytes (CD14-CD16+) utilize OXPHOS. Relevant to MS, Fitzgerald et al. reported significant changes in circulating aromatic amino acid metabolites in MS, which correlated with disease activity in people with MS^39^. Their study also found reduction in amino acid metabolites within CD14+ monocytes as well as in CSF and blood, which in turn was found to be associated with increased disease activity in people with MS. While alterations in amino acids are influenced by many factors such as gut microbiota, metabolic pathways within immune cells such as glycolysis and TCA cycle are directly tied to immune activation profiles through amino acid metabolism. Moreover, amino acids directly influence OXPHOS as amino acid metabolites act as precursors to TCA cycle intermediates^40^. Thus, reduction in amino acid metabolites likely forces monocytes to switch to glycolysis to compensate for the loss of ATP, and to assist with proinflammatory functions.

Our previous studies showed that CSF samples from SPMS patients had elevated soluble EMMPRIN levels^13^, while CNS-infiltrating monocyte-derived macrophages increased their EMMPRIN expression at sites of infiltration in MS brains^9^. Our study also established a strong link between EMMPRIN and MCT-4, assisting with lactate secretion, in mediating pro-inflammatory activation of macrophages in EAE^9^. In the current study, using newly generated EMMPRIN floxed mice, we specifically examined EMMPRIN with regards to its roles in CCR2+ monocytes before they manifest as tissue-infiltrating macrophages in EAE pathology. Our findings, for the first time, provide evidence for EMMPRIN in assisting specific metabolic pathways within CCR2+ monocytes during EAE. The majority of the pathways in CCR2:EMMP^−/−^ BMDMs exhibited increase in fatty acid oxidation and mitochondrial respiration and decrease in glycolysis, which is associated with anti-inflammatory or homeostatic functions in macrophages. scRNA-seq datasets of blood monocytes from CCR2:EMMP^−/−^ mice revealed more complicated phenotypes since EMMPRIN knockdown downregulated genes pertaining to glycolysis and mitochondrial respiration in D7 EAE monocytes. It is likely that inherent differences exist in the metabolic pathways between circulating monocytes and tissue resident macrophages. It is also noteworthy that during chronic inflammation, macrophages reprogram to immunosuppressive phenotypes by reducing their mitochondrial translation^36^. As seen in dendritic cells, an increase in mitochondrial activity is linked to higher ROS production, which is mitigated by elevated intracellular lactate levels through the HIF-1α pathway^41^. This observation implies that blocking lactate export may induce anti-inflammatory responses in myeloid cells, potentially explaining the effects seen with EMMPRIN deletion in our study. Recently, another study showed reverse electron transport through mitochondrial complex I to induce pathological phenotypes in microglia^33^. Thus, the observation from scRNA-seq experiment that mitochondrial respiration-associated genes were downregulated within CCR2:EMMP^−/−^ monocytes point to the pro-inflammatory nature of EMMPRIN^36^. An earlier study showed that THP-1 human monocytes underwent significant metabolic changes in response to inflammation where early activation enhanced glycolytic pathway intermediates such as glucose 1-phosphate and fructose 1, 6-bisphosphate^42^. Further, fatty acid synthesis, a phenotype opposite to fatty acid oxidation, also enhanced in these monocytes during inflammation^42^. These observations confirm that monocytes bear complicated metabolic phenotypes which are likely governed by chronicity of inflammation. At the same time, it cannot be refuted that some of the differences with respect to the metabolic pathways in monocytes and macrophages could be due to the comparison between in vivo and in vitro systems. Overall, our collective findings implicate EMMPRIN as a key regulator of metabolic pathways that exacerbate pro-inflammatory functions of monocytes and macrophages in MS.

We were surprised that while EMMPRIN was initially described as an inducer of MMPs ^31, 43^, levels of MMPs were not altered in transcripts or protein content in the scRNA-seq or proteomic analyses, respectively. This emphasizes the many roles of EMMPRIN^44^ that have been described in addition to the capacity of EMMPRIN in other context to induce MMP expression. It is also possible that the MMP inducing roles of EMMPRIN reside in non-CCR2 immune cell types. Lack of EMMPRIN’s interaction with its cognate ligands such as soluble cyclophilins, which might be required for MMP production in monocytes^32^, might also contribute to these observations. These possibilities deserve examination in future studies.

Our research aligns with observations indicating the presence of distinct subgroups of monocytes during EAE pathology^45^. In this regard, identification of the altered metabolic programs within individual subclusters in CCR2:EMMP^−/−^ mice would be informative. However, due to constraints in the number of cells from this study, we were unable to decipher meaningful differences in metabolic pathways within each subcluster. Nevertheless, EMMPRIN deletion in CCR2:EMMP^−/−^ mice had similar outcomes in EAE as to inhibiting MCT-4 in macrophages^9^, which is likely due to reduction of transmigration of CCR2:EMMP^−/−^ monocytes to the CNS. Another important finding of the current study is the disruption of differentiation-assisting genetic make-up. For instance, we found increase in a myelopoiesis assisting gene, *Slfn4,* in CCR2:EMMP^−/−^ Cluster 8 which is associated with myeloid progenitor cells^24^. In this regard, we confirmed a decrease in Slfn4 in BMDMs, which correlated with enhanced EMMPRIN levels during inflammation. These findings are in alignment with previous observations that reducing EMMPRIN in monocytes interfered with their differentiation program^43^. Additionally, enhancement in transcripts of *Csf3r* and *Ly6a*, which are associated with stemness of progenitor cells, were elevated in CCR2:EMMP^−/−^ monocytes. This indicates that EMMPRIN depletion potentially prevented pro-inflammatory monocytes from differentiating to tissue-resident macrophages during EAE pathology. This might explain why the majority of monocyte subclusters were depleted in CCR2:EMMP^−/−^ mice and potentially why significantly less macrophages transmigrated the EAE CNS. This interpretation is bolstered by the observations that CD115+ (CSF1R) monocytes were significantly reduced in CCR2:EMMP^−/−^ mice in D7 EAE when compared with the controls.

Intriguingly, elevation of CSF-1 in CCR2:EMMP^−/−^ macrophages *in vitro* signifies compensatory mechanisms to override differentiation-inhibiting pathways in these macrophages. Observations regarding decrease in *slc16a3* in CCR2:EMMP^−/−^ BMDMs were intriguing. While EMMPRIN chaperones MCT-4 to the macrophage membranes and that SiRNA-mediated knockdown of EMMPRIN reduces membrane placement of MCT-4^9^, EMMPRIN’s role in transcriptional regulation of MCT-4 was not described earlier. It is noteworthy that CCR2:EMMP^−/−^ mice did not reduce circulating Ly6G-CD45+ CD11b+ monocytes in the blood of non-EAE healthy mice. This suggests that directing therapeutic efforts towards EMMPRIN holds great potential, as it targets pro-inflammatory monocytes and monocyte-derived macrophages specifically during EAE.

In studies of active lesions in autopsied brain specimens of MS, it has been emphasized that myeloid cells (monocyte-derived macrophages and microglia) constitute over 90% of the immune cell repertoire^46, 47^. Positron emission tomography of living individuals show that the density of myeloid cells is increased from early disease and are particularly elevated in those with progressive MS^48, 49^. Where immunostains are available to define the cell subsets in autopsied lesions, macrophages (CD68+TMEM119-) account for ∼70% of myeloid cells in active lesions, and ∼50% at the rim of chronic active lesions^50^. Thus, besides an increasing interest in affecting microglia in MS, it is important to restore homeostasis of monocytes and macrophages in MS. Our results would suggest that affecting EMMPRIN biology would be one means to normalize the metabolism of monocytes and macrophages, leading to reduced disease activity in MS.

## ACKNOWLEDGMENT

This work was supported by a Foundation grant from the Canadian Institutes of Health Research (VWY) and by an Exploration – Hypothesis Development grant from the Congressionally Directed Multiple Sclerosis program of the USA Department of Defense (DKK) and MS Canada (DKK). We thank Ken Ito and Derek Rancourt of the Centre for Genomic Engineering of the Cumming School of Medicine for deriving the EMMPRIN^fl/fl^ mice. We thank Nick Newhook and Stephanie Tucker at Medical Laboratories (Memorial University) for their technical assistance. We thank Dr. Lydia Sorokin (University of Münster, Germany) for the kind gift of rabbit anti-pan laminin antibody.

## AUTHOR CONTRIBUTIONS

D.K.K. and V.W.Y conceived the study, designed the project and co-supervised the overall study. V.W.Y and M.X. discussed various aspects of EMMPRIN biology and the necessity of generating EMMPRIN^fl/fl^ mice. D.K.K characterized the EMMPRIN^fl/fl^ mice, performed the experiments and analyzed data. E.P.N. and N.J.B. provided critical support with 10X Genomics for scRNA-seq. C.D.M. analyzed the single cell dataset and assisted with writing the associated results. A.D. (Univ. of Calgary) and L.A. conducted LC-MS/MS analyses. C.S., A.D. (Memorial University) and N.G. maintained the CCR2Cre^ERT2^:EMMPRIN^fl/fl^ lines and helped with macrophage differentiation experiments. D.K.K. wrote the first draft of the manuscript. All authors reviewed and approved the manuscript.

## MATERIALS AND METHODS

### Generation and characterization of CCR2:EMMP^−/−^ mice

EMMPRIN^fl/fl^ mice were generated by inserting flox sequences flanking the 5’ region of exon 1 and 3’ region of exon 8 of *Bsg* (Basigin), the EMMPRIN gene (Centre for Genomics Engineering, Cumming School of Medicine, University of Calgary; Supplemental Fig 1). LoxP sequences were inserted by microinjecting CRISPR/Cas9 and gRNA constructs in oocytes from DBA2 strain and the insertion was confirmed by gene sequencing. The mice were backcrossed into C57BL/6 strain for 6 generations to attain >95% purity on C57BL/6 background. The following genotyping primers were used to confirm flox insertion in the *Bsg* gene: 5’ forward primer: 5’- GGGTTCTTGGGGTAAAGAGG-3’, 5’ reverse primer: 5’-AGTCACATGGCCCACTTCTC-3’, 3’ forward primer: 5’-AGAAGCCTAGGGCAACCACT-3’ and 3’ reverse primer: 5’- CTTCGCCAGCCTGTCTTACT-3’.

The CCR2Cre^ERT2-^mKATE mice were procured from Dr. Burkhard Becher^6^ and crossed with EMMPRIN^fl/fl^ mice to generate the tamoxifen-inducible CCR2Cre^ERT2^:EMMPRIN^fl/fl^ (CCR2:EMMP) mice. CCR2:EMMP, EMMPRIN^fl/fl^ and CCR2Cre^ERT2^ lines were maintained as homozygous. EMMPRIN^fl/fl^ and CCR2:EMMP^−/−^ mice were 100% viable and exhibited no obvious developmental or any other health issues. CCR2Cre^ERT2^ had high recombination efficiency as homozygous as previously published^6^, which we confirmed using the Ai9-Cre reporter mice (The Jackson Laboratory; strain # 007909) (Supplementary Fig 2). For CCR2Cre^ERT2^, the following primers were used for genotyping: forward primer: 5’-AGAAAGTGAGCCCTCTGTATGG-3’ and reverse primer: 5’-TTGGCATTTCCTGGTGAGC-3’. All the animal experiments were conducted in accordance with Canadian Council on Animal Care guidelines and with ethics approval from the University Animal Care Committee at University of Calgary, Canada and Memorial University of Newfoundland, Canada.

### EAE induction, tam-injections, confocal imaging, and flow cytometry

We induced EAE in CCR2:EMMP^−/−^ homozygous mice by subcutaneous injections of 50µg of myelin oligodendrocyte glycoprotein (MOG)35-55 peptide (Protein and nucleic acid facility, Stanford, CA) emulsified in complete Freund’s adjuvant (fortified with heat-inactivated 10mg/ml *M. tuberculosis* H37RA (Thermo Fisher Scientific, Toronto, Canada) in 6-8 weeks old male and female CCR2:EMMP^−/−^ mice. Tam (T5648, Sigma) was reconstituted in corn oil and injected at 2mg/mouse subcutaneously in l00 μL volume. Animals were monitored daily for clinical signs of EAE on the 15-point scale ^9,51^.

For immunohistochemistry, EAE and control mice were anesthetized with a lethal dose of ketamine and xylazine and perfused with ice-cold PBS. Cerebellums were then harvested, fixed in OCT medium (23-730-571, Fisher Scientific), sectioned at 20- to 30-μm thickness, and stored at −80°C until further processing. Cerebellum sections were then fixed in 100% methanol at –20°C and incubated with rat anti–mouse CD45 (550539, BD Pharmingen, 1:100) and rabbit anti–pan laminin (a gift from L. Sorokin, University of Münster, Münster, Germany; 1:1000) antibodies for staining of perivascular cuffs. After additional washes, sections were incubated with Alexa Fluor 488 and Alexa Fluor 546 secondary antibodies (Invitrogen, 1:500) and mounted with Fluoromount-G^TM^ media containing DAPI (00-4959-52, Invitrogen). Images were captured on a Fluoview FV10i confocal microscope (Olympus) and AiryScan Confocal microscope (Zeiss, Memorial University).

For flow cytometry, approx. 0.5mL heparinized whole blood was collected using cardiac puncture from EAE (D7, D12) and control mice and subjected to RBC lysis using RBC lysis solution (555899, BD Biosciences) before proceeding with flow cytometry. For spinal cord flow cytometry, spinal cord tissues were homogenized following transcardial perfusion using ice cold 1X PBS and subjected to myelin removal using the Percoll gradient method. The interphase was then re-suspended in FACS buffer and blocked with the Fc-blocker CD16/CD32 (553141, BD Biosciences, 1:100) for 20 minutes at 4°C. Cells were then washed and incubated in following antibodies for 30 minutes at 4°C in the dark: PerCP rat anti-mouse CD45 (557235, BD Biosciences, 1:50), FITC rat anti-CD11b (553310, BD Biosciences, 1:50), APC-Cy7 rat anti-mouse Ly6G (560600, BD Biosciences, 1:50), PE rat anti-mouse EMMPRIN (562676, BD Biosciences, 1:50), BUV395 rat anti-mouse CD115 (564059, BD Biosciences, 1:50), BV510 rat anti-mouse CCR2 (CD192, 747970, BD Biosciences, 1:50) and fixable viability dye eFluor 780 (65-0865-14, eBioscience). Cells were then washed in FACS buffer and fixed in 1% formalin before being suspended in FACS buffer and analyzed using a BD LSR II FACS system and Cytoflex (Beckman Coulter, Memorial University). Cell populations were gated based on viability dye and subjected to forward and side scattering to identify single cell population before additional analysis using Flowjo (BD Biosciences).

### BMDM isolation, Luminex assay, qPCR analysis, Lactate measurement and Western blotting

Bone marrow from CCR2:EMMP^−/−^ mice and WT mice were flushed out, pelleted at 1,200 rpm for 10 mins and resuspended in complete high glucose DMEM media (10% FBS, 2% penicillin/streptomycin) supplemented with 10% supernatant from the L929 fibroblast cell-line. The cells were seeded at a density of 10^7^ on 10cm petri dishes as previously described ^52^ and incubated at 37°C in 8.5% CO_2_ for 5 days, after which half of the medium was replaced with fresh medium. At D7, the full medium was replaced with fresh medium, and the cells were seeded in 6-well plates and 96-well plates for various assays including Western blotting, qRT-PCR, LC-MS/MS and Luminex analysis. The cells were treated with 5µM 4-OH tam (H6278, Sigma) for 24h to delete EMMPRIN in tam and LPS+tam conditions. For LPS treatments, BMDMs were stimulated with 100ng/ml LPS for 24h before subjecting to downstream applications. For cytokine analysis, cell culture supernatant from BMDMs was subjected to 31-plex chemokine/cytokine luminex assay (Eve Technologies, MD31).

For Quantitative PCR, BMDMs were lyzed using the Trizol method (15596026, Invitrogen) and RNA was harvested using the RNAeasy Kit (74104, Qiagen). The isolated RNA from different conditions was subjected to qPCR using the Fast SYBR green master mix (4385612, Applied Biosystems) on a Quantstudio 6 Flex (Applied Biosystems). The following primers were used: Silencer select^tm^ (ThermoFisher Scientific) primers for *hk1* (assay ID. 758971) and *slc16a3* (assay ID. 174958) and *Actb* (QIAGEN ref. QT00095242; NCBI RefSeq: NM_007393). We generated the following primers for EMMPRIN (*Bsg*): forward primer: 5’- GCGGCGGGCACCATCCAAAC-3’; reverse primer: 5’ ATGTACTTCGTATGCAGGTCGG-3’. Transcripts were analyzed using the ΔΔCt method, and the data were normalized to *Actb*.

For lactate measurement, cell culture supernatant was analyzed for lactate levels using the fluorescence-based L-lactate measurement kit according to the manufacturer’s protocol (ab65330, Abcam). Briefly, cell culture supernatants (1:200 primary dilution) and the standards were incubated in the presence of lactate probes and enzyme mix at room temperature for 30 minutes and measured at 535/587 nm excitation/emission using Spectramax Mini plate reader (Molecular Devices). Lactate levels were represented in millimolar concentrations.

For Western blot analysis, BMDMs were lysed in RIPA buffer (89900, ThermoFisher Scientific) supplemented with protease and phosphatase inhibitor (78440, ThermoFisher Scientific) and 30 μg of BMDM lysates were loaded onto a 10% polyacrylamide gel and transferred onto PVDF membranes (Bio-Rad Laboratories). The membranes were then blocked with 5% non-fat milk for 1.5h and incubated with rabbit anti-Slfn2/3/4 (PAB5934, Abnova, 1:1000), rat anti-EMMPRIN (NB100-65518, Novus Biologicals, 1:1000), rabbit anti-EMMPRIN (Ab108317, Abcam, 1:1000, used to confirm knockdown), and HRP conjugated β-actin (ab20272, Abcam, 1:5000) antibodies overnight at 4°C in 1% skim milk. The following day, membranes were washed with 0.1% 1x TBS Tween-20 (TBST) and incubated with HRP-linked anti-rabbit (A16023, Invitrogen, 1:5000) or anti-rat (A18739, Invitrogen, 1:5000) antibodies for 1.5h in TBST and then washed before chemiluminescence imaging (Bio-Rad Laboratories).

### CD11b+ cell isolation and scRNA-seq

For scRNA seq analysis, we collected peripheral blood from 3 CCR2:EMMP^−/−^ and 3 CCR2:EMMP^+/+^ D7 EAE mice that received tamoxifen via transcardial perfusion. CD11b+ cell isolation kit (StemCell CD11b+ positive selection kit; Cat#18970A) was then used to magnetically sort CD11b+ cells. Briefly, blood was subjected to RBC lysis and 1X10^8^ cells per ml were resuspended per ml of MACS buffer provided in the kit. The cells were then added with 12.5µl of rat serum and moved to polystyrene tubes to fit the Easy Sep magnets. 12.5µl of a mixture of component A+B was then added to the mix and incubated at room temperature (RT) for 5 min. Following this, 20µL of rapid spheres were added to the samples and incubated for 3 min at RT. Using MACS buffer the solution was topped up to 2.5ml and the tubes were placed on magnetic stand for 3 min at RT. This resulted in >97% viability of CD11b+ cells. Target recovery/sample was set at 5000 cells and planned sequencing platform for 800M reads. We then performed scRNA-seq using chromium next GEM single cell kits^tm^ from 10x Genomics followed by sequencing using NovaSeq Ilumina system. Cell ranger pipeline recovered 3001, 2714 and 3608 cells for the no-tam ‘mice and 3560, 3855 and 3796 cells for the 3 CCR2:EMMP^−/−^ group with most reads >30,000 reads/cell. After processing the scRNA-seq data by Cell Ranger pipeline, the expression matrix was analyzed using the Seurat package v.3 in R to identify major metabolic pathways in monocytes identified by Gene ontology and Ingenuity Pathway Analysis (IPA).

### Single Cell 3’ RNA Library Construction

The CD11b+ cells obtained post sorting were processed with the 10x Genomics Chromium single cell platform. Appropriate volume of cells to aim for a target recovery of 5000 cells/library was loaded on the Chromium controller chip for single cell partitioning and barcoding. Post cDNA amplification, library construction was performed as per the Chromium single cell 3’ reagent kits user guide v3.1. cDNA and library construction quality control were performed using the Agilent Tape station. All 6 libraries were sequenced on the Novaseq Illumina platform using the SP flowcell with run parameters as follows: 28 cycles read 1, 10 cycles for the i5 and i7 index and 90 cycles for read 2.

### Single Cell RNA sequencing data analysis

Sequencing files were demultiplexed using the 10x Genomics Cell Ranger version 5.0 pipeline. Briefly, FastQ files were processed with the Cell Ranger count pipeline which uses STAR to align reads to the mouse reference genome mm10. The Cell Ranger aggregate pipeline was run to generate an expression matrix with the 6 combined libraries with normalization setting set to ‘None’. The final expression matrix with gene counts were then analysed using the bioinformatics package Seurat v3 in the R programming environment. Prior to running QC metrics, the expression matrix comprised 16,382 features and 20,518 cells. The data was filtered for the following parameters: gene present in > 3 cells, cells with < 6000 genes and percent of mitochondrial genes < 20%. Post filtering, the expression matrix contained 16,382 features and 20,306 cells. Data was normalized and scaled using the SCTransform function with nCount_RNA and percent.mt being regressed out. A PCA-reduction was performed, and 30 significant PCA-dimensions were considered. Clusters were determined using the FindClusters function which implements the Louvain algorithm for modularity optimization and was performed with a resolution of 0.5. Cluster annotation was done manually based on the expression of lineage specific hallmark genes. DEGs for one cluster (versus all cells in other clusters) was determined by the default Wilcoxon rank sum test. To assess monocyte subsets, the monocyte cluster was subset from all other cells and re-clustered using 20 significant PCA-dimensions and 0.5 resolution. To determine enrichment of functional pathways, significant DEGs were imported into IPA software and EnrichR gene enrichment tools.

### Data availability

Single cell transcriptomic data will be deposited on the NCBI Sequence Read Archive (SRA) site and accession code made available accordingly.

### Proteomics analysis

For shotgun proteomics of cell culture samples, the cells upon treatment with 24h 4-OH followed by 24h LPS stimulation were pelleted and lysed in 1% SDS, 100mM ammonium bicarbonate, 10mM EDTA, and 1 cOmplete™ tablet (Roche, 4693159001), for a final pH of 8. Shearing of the DNA was accomplished by sonicating the samples on ice, followed by centrifugation (10,000g, 10 min, 4 °C) and transfer of the supernatant to a new tube. A total protein amount of 100µg per sample was incubated with a final concentration of 20% trichloroacetic acid (TCA) on ice for 30 min, followed by centrifugation (14,000 x *g*, 15 min, 4 °C) for precipitation. The pellet was washed three times with cold acetone (4 °C). The final pellet was left to air dry (2 min) and resuspended with 8M urea in 0.1 M Tris-HCl (urea-tris), pH 8. Samples were digested with trypsin using the filter aided sample preparation (FASP) method^53^. Peptides were eluted with 50 mM of ABC, lyophilized, and desalted in the Pierce™ Peptide Desalting Spin Columns (89851, ThermoFisher Scientific). Prior to labelling, peptides were lyophilized and resuspended in 100µL of 50mM triethylammonium bicarbonate (TEAB), pH 8. Peptides were labelled using the TMT-6plex™ Isobaric Labeling (90061, ThermoFisher Scientific) according to the manufacturer’s directions. Briefly, the TMT label reagents were resuspended in anhydrous acetonitrile and added to each sample (41µL TMT 6plex™ per 100 µL sample) and incubated at RT for 1h. The TMT labeling reaction was quenched by 8 µL of 5% hydroxylamine for 15 min at RT and the samples were combined. Samples were stored at −80 °C before lyophilization, followed by resuspension in 1% formic acid before liquid chromatography and tandem mass spectrometry analysis for which Peptides were analyzed on an Orbitrap Fusion Lumos Tribrid mass spectrometer (ThermoFisher Scientific) combined to a Thermo Scientific Easy-nLC (nanoflow liquid chromatography) 1200 System and operated with Xcalibur (version 4.0.21.10).

### Proteomic data and bioinformatics analysis

Spectral data acquired from the mass spectrometer were matched to peptide sequences in the murine proteome from the UniProt database using the Andromeda algorithm^54^ in the MaxQuant^55^ software (v.1.6.0.1), at a peptide-spectrum match false discovery rate (FDR) of 0.01. The normalization and identification of differentially expressed proteins was performed using the MSstatsTMT package^56^. Multiple comparisons were corrected using the Benjamini-Hochberg approach. For protein-protein interaction analysis, proteins with an adjusted p-value <0.05 and a log fold change >1 or <-1 were submitted to the STRING (Search Tool for the Retrieval of Interacting Genes) database^57^. The same set of proteins were submitted to Metascape^58^ as a “Multiple Gene List” for pathway enrichment analysis and represented as heatmaps. Custom heatmaps were plotted using the heatmap.2 function from the gplots package^9^. The Venn diagram was plotted using the VennDiagram package. Volcano plots were plotted using the ggplot2 package. The proteomics data are publicly available and are deposited in PRIDE Archives. The R codes are available upon request.

