## Supplementary figures and images for "EMMPRIN confers metabolic advantage for monocytes and macrophages to promote disease in a model of multiple sclerosis"

### Supplemental Fig S1

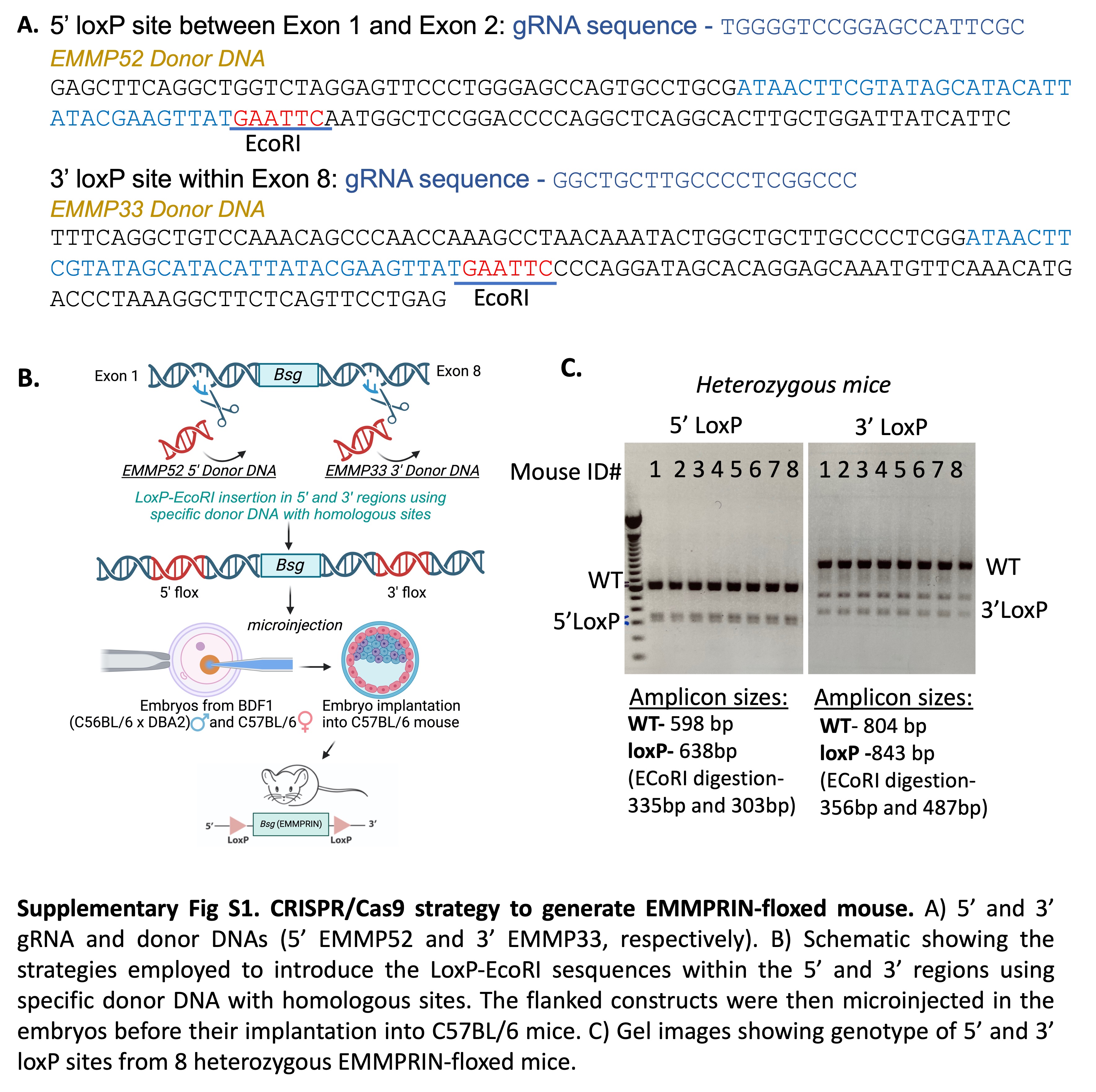

### Supplemental Fig S2

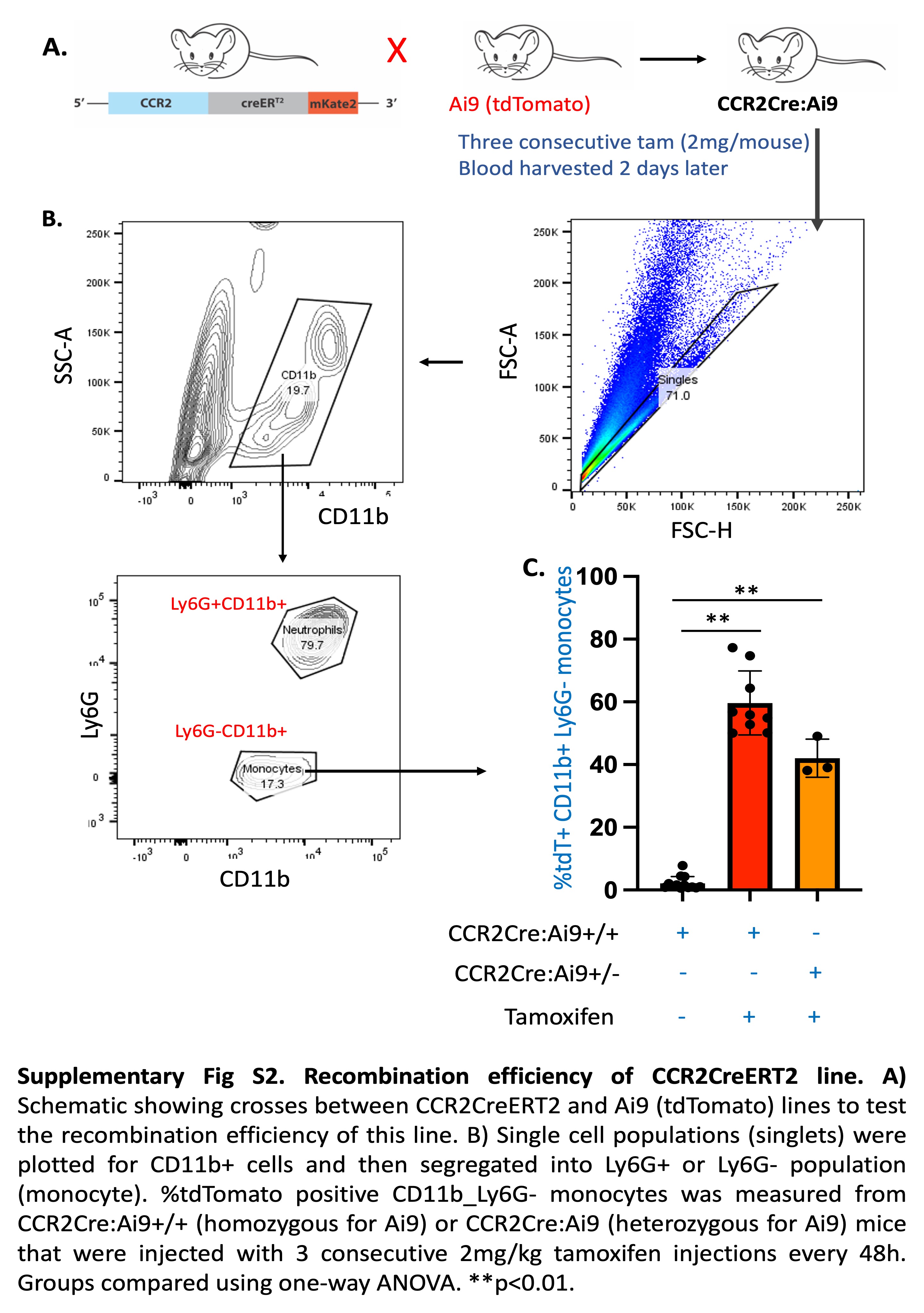

### Supplemental Fig S3

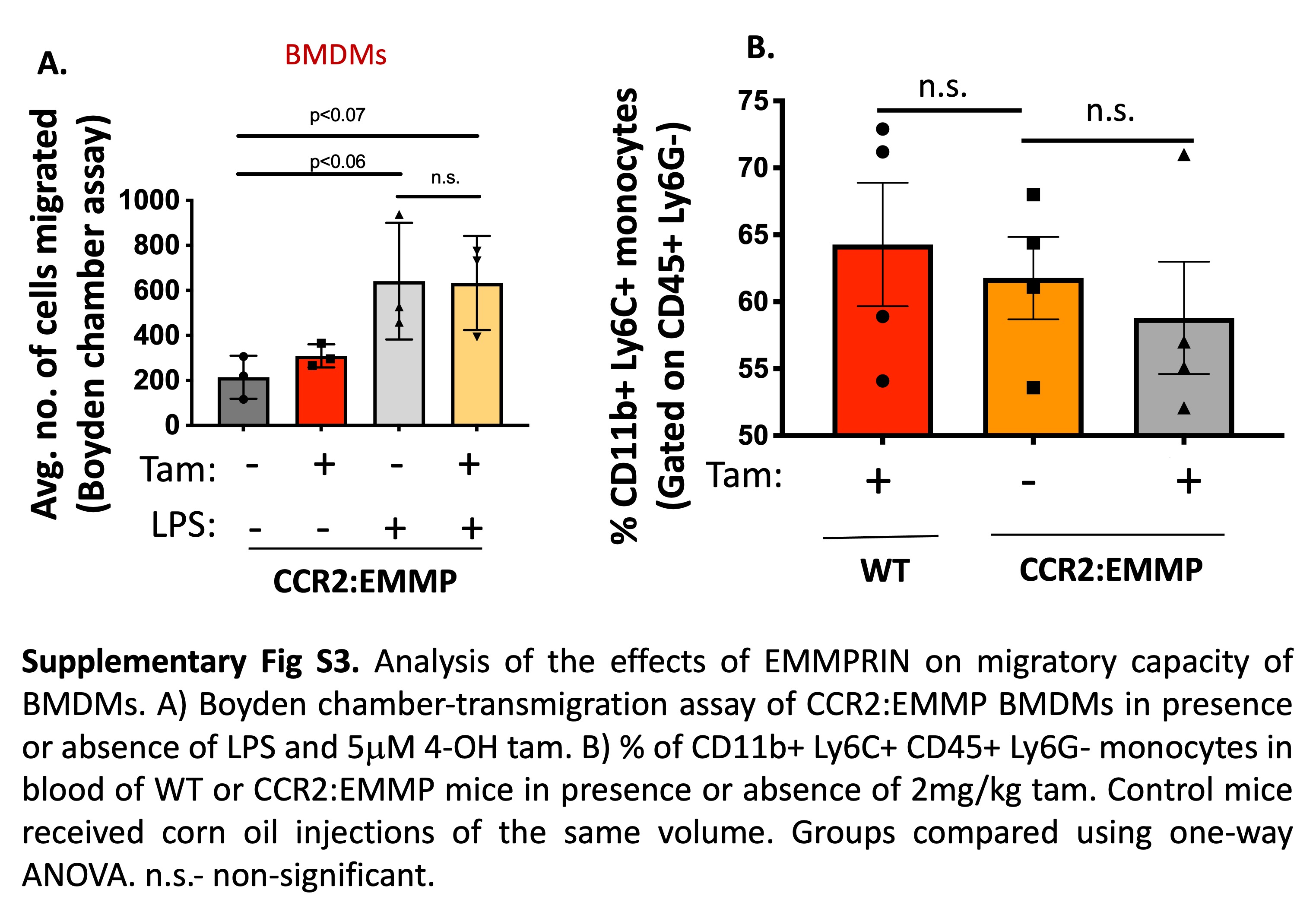

### Supplemental Fig S4

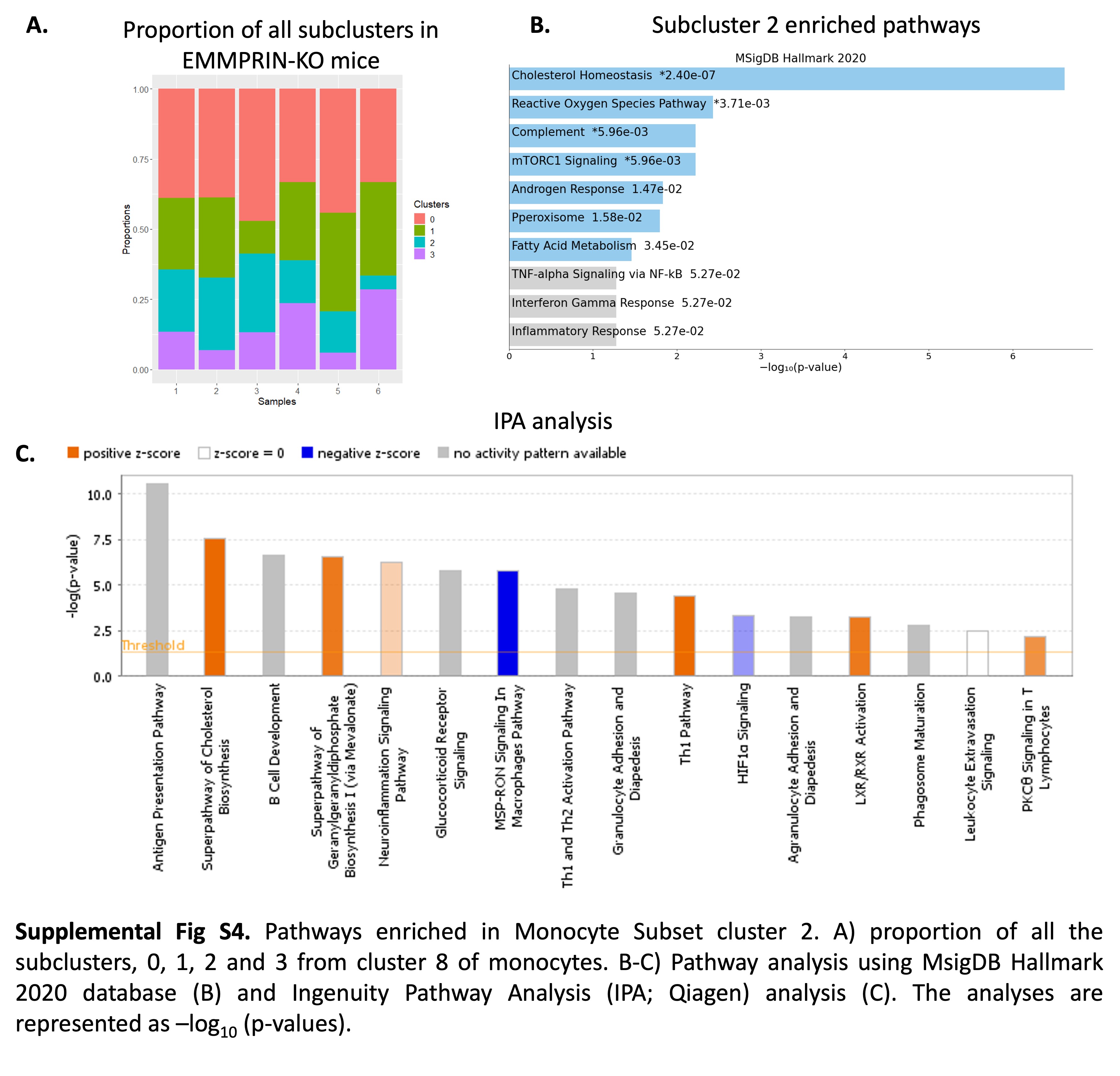

### Supplemental Fig S5

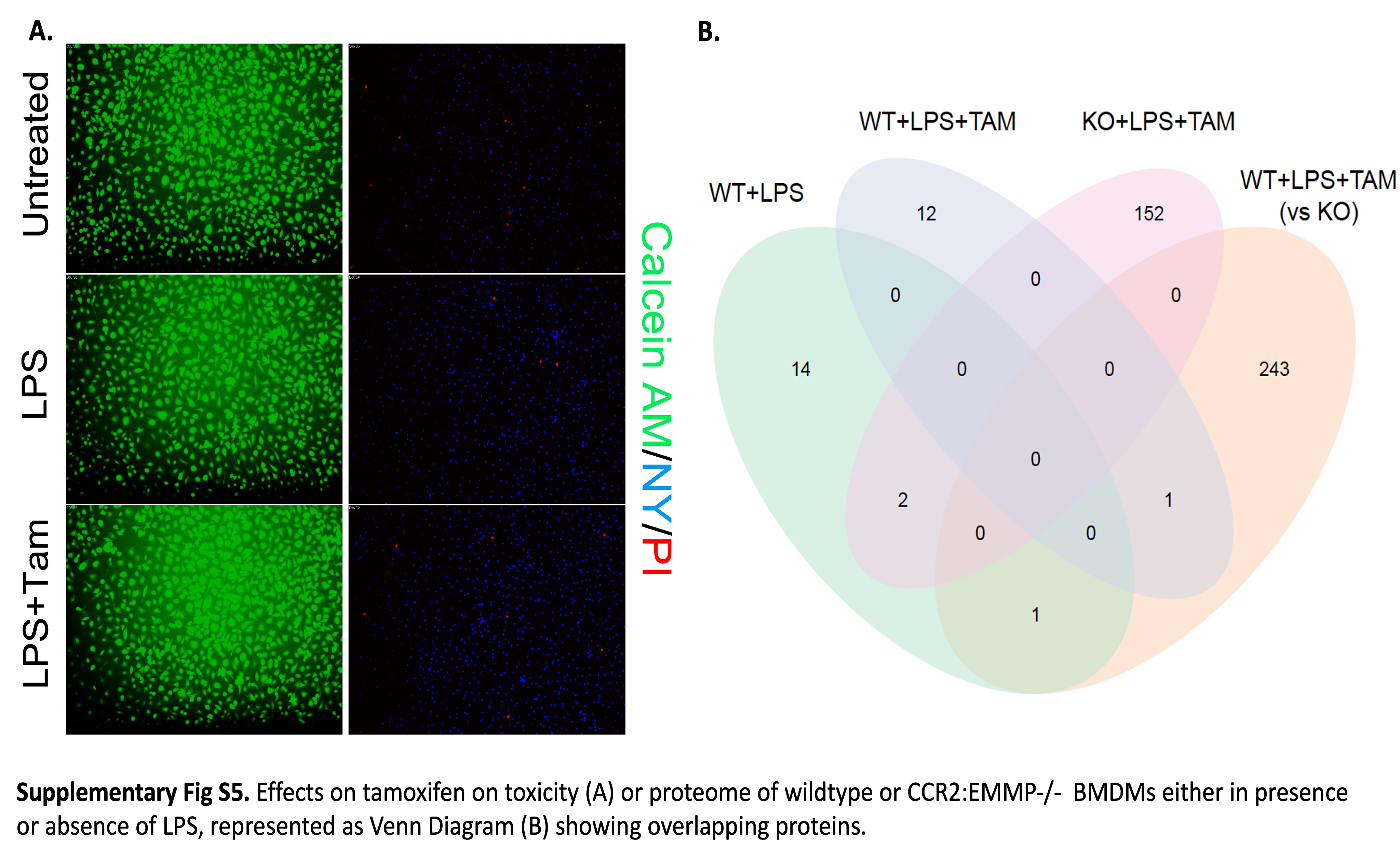
